# Sensitivity of Gene Sets to miRNA Regulation: A Cell-Based Probabilistic Approach

**DOI:** 10.1101/2020.05.09.778589

**Authors:** Shelly Mahlab-Aviv, Nathan Linial, Michal Linial

## Abstract

Mature microRNAs (miRNAs) are small, non-coding RNA molecules that function by base-pairing with mRNAs. In multicellular organisms, miRNAs lead to mRNA destabilization and translation arrest. Importantly, the quantities and stichometry of miRNAs/mRNAs determine the miRNA regulation characteristics of specific cells. In this study, we used COMICS (Competition of miRNA Interactions in Cell Systems), a stochastic computational iterative framework to characterize genes by their sensitivity and robustness to miRNA regulation. We monitor the cell state by quantifying the retention level for all mRNAs, at the end of 100,000 simulation iterations. In HeLa cells, we partitioned all genes to five classes according to their decay rates. We show that the largest class (69% of genes) is apparently resistant to miRNA regulation. We created in silico perturbations using overexpressing of all major miRNAs (248 types) at various levels relative to the basal level (x1 to x1000). We further classified genes according to the differential behaviour for any pair of conditions of miRNA expression profile. Based on such measure (OvereXpression Ratio, OXR), we identified a small number of gene sets that are especially sensitive to OXR. Our results expose an overlooked quantitative dimension for set of genes and miRNA regulation in living cells.

## 1 INTRODUCTION

Mature microRNAs (miRNAs) are non-coding RNA molecules (∼22 nucleotides) that regulate genes through base complementarity with their cognate mRNAs, at the 3’-untranslated regions (3’-UTR) (Moore et al., 2015). In multicellular organisms, miRNAs act by destabilization of mRNAs and interfering with the translation machinery (Chekulaeva and Filipowicz, 2009; Eichhorn et al., 2014). Alteration in the relative abundance of miRNAs may lead to transition between states, and the establishment of cell identity (Pelaez and Carthew, 2012). Changes in miRNA cell composition is associated with viral infection (Zhang et al., 2012), differentiation, and cancer transformation (Bertoli et al., 2015; Lu et al., 2005).

The current catalogue of miRNAs (Kozomara and Griffiths-Jones, 2013) contains ∼2500 mature miRNAs from humans. However, the number of miRNA types that are expressed at a substantial amount in most cells is much smaller. The miRNA regulatory network can be formulated as a bipartite graph with nodes at one side correspond to gene transcripts, and on the other side to miRNAs. Many miRNAs carry the potential for targeting hundreds of transcripts (Balaga et al., 2012; Rajewsky, 2006), and tens of miRNA binding sites (MBS) are predicted at the 3’-UTR of mRNAs (Landgraf et al., 2007). Most of our current knowledge on the specificity of miRNA-mRNA regulatory network is based on computational prediction tools (Peterson et al., 2014) that suffer from a flood of false positives (Pinzon et al., 2017). Results from CLIP-based protocols provide rich, unpaired collection of miRNAs and mRNAs from living cells (Li et al., 2014). Unfortunately, many of the deep sequencing-based protocols suffer from low coverage and poor consistency (discussed in (Lu and Leslie, 2016)). It is expected that poorly expressed miRNAs do not contribute to downregulation of gene expression (Hausser and Zavolan, 2014). In reality, only ∼60% of the human coding genes are estimated for being regulated by miRNAs in vivo (Ha and Kim, 2014; Jonas and Izaurralde, 2015).

The properties of the miRNA-mRNA regulation in a specific cell type depend on the amounts and concentration of miRNAs and their stoichiometry. A key player in the regulation is the availability of AGO protein, the catalytic component of the RNA silencing complex (RISC) (Janas et al., 2012; Wen et al., 2011). In view of mRNAs, the abundance and positions of MBS along the relevant transcript (Jens and Rajewsky, 2015) dictate the potential of miRNA interaction, but not necessarily the potency for gene expression attenuation (Agarwal et al., 2015). The properties of the miRNA-mRNA complex network call for an unbiased probabilistic model for defining the principle design of miRNA regulation.

In this paper, we develop a quantitative view on miRNA regulation. We provide a stochastic - probabilistic model that operates at the cellular level. We configure an iterative simulator on cell-line by an exhaustive set of miRNA manipulations by an overexpression paradigm. In the end of the simulation run the mRNA retention levels are monitored and a new steady state is defined. We created two unbiased gene sets - the first one concerns the dynamic behaviour along the simulation run (dynamic-classes). The second classification concerns gene class by their sensitivity to changes in miRNA quantities, based on the end-point retention ratio for any pairs of overexpression settings, these are called OXR-classes (OvereXpression Ratio-classes). The rich representations by the dynamics-and OXR-classes expose overlooked properties of miRNA regulation.

## 2 METHODS

### 2.1 Probabilistic map for miRNA-mRNA pairing

The probabilistic framework interaction table used in this study was adapted from the scores provided by TargetScan (Agarwal et al., 2015). Accordingly, high probability of successful interactions is calculated from a combination of strongly supported miRNA-mRNA pairs that comply with many features from sequence, secondary structure and evolution conservation. The complete miRNA-mRNA interaction table includes 8.22 M pairs covering the highly and poorly conserved interactions. We used a compressed version of the table that reports only on pairs that are supported by evolutionary conserved miRNAs with total of 1,183,166 pairs, covering 18,953 genes and 289 miRNA families. The predicted repression scores range from 0-1, and are identical for all representation of the relevant miRNA family members (Agarwal et al., 2015). The conversion of the interaction scores to binding probabilities was done according to TargetScan score: p = 1 – 2^score.

#### 2.2.1 Normalizations of mRNA expression and miRNA families

For mRNA expression profile, we used experimental data that we have produced (HeLa cell, expression by mapped reads (Mahlab-Aviv, 2018). We reliably mapped 16,355 mRNAs and 539 miRNAs that were expressed at a minimal level of >=1 reads (from total mapped reads ∼3 M).

For miRNA normalization we estimated 50,000 molecules per cell and miRNAs with more than 1 molecule after normalization was considered (total 110 miRNAs). Similarly, we estimated 25,000 mRNA molecules, and following the expression threshold (total 3666 mRNAs). For the generic miRNA list, miRNAs are compiled to match the definitions provided by the TargetScan interaction matrix, a total of 248 miRNAs according to TargetScan table. The analysis that are based on % retention. The analysis is bounded by genes having a minimal number of molecules (e.g., 0.02% of cell expression, 5 mRNA molecules, total 753 genes).

### 2.2 Probabilistic based miRNA-mRNA simulator

The simulator inputs include the number of molecules from the expression profiles of miRNAs (50k molecules) and mRNAs (25k molecules), and a table of miRNA-mRNA interaction prediction extracted from TargetScan. In each run, a random miRNA is chosen from the predetermined available miRNAs distribution. Next, a target is chosen randomly according to the available targets’ distribution. mRNA that is already bounded by other miRNA molecules can still be a putative target for the chosen miRNA, if the relevant binding site is does not overlap with an occupied MBS on the same molecule. Overlapping binding sites are considered any neighbouring MBS which are <50 nucleotides apart. A binding event will occur according to the estimated miRNA-mRNA binding probability from the TargetScan interaction table. Upon a binding event, the free miRNA and mRNA distributions are updated, and the bounded mRNA molecules are marked as being occupied. An occupied molecule is removed after 1k iterations following a successful binding event. For mRNA to be eliminated, at least one MBS must be reported as occupied. After mRNA removal, the bound miRNAs are released and return to the free miRNAs. Thus, become eligible for ongoing binding events.

#### 2.2.1 Configuration of COMICS

COMICS (Competition of miRNA Interactions in Cell Systems) simulator supports a wide set of configurable parameters: (i) the number of total miRNA; (ii) the number of mRNA molecules in the cell; (iii) the number of iterations for completing the run; (iv) the number of iteration interval between miRNA-mRNA binding event and the mRNA removal; (v) a random removal of unbounded mRNAs according to predetermined decay rate of the mRNA as extrapolated from experimental mRNA half-life; (vi) addition of newly transcribed mRNAs along the iterations interval; (vii) miRNAs or genes overexpressed according to a selected multiplication factor; (vii) incorporation of alternative miRNA-target mapping. It is also possible to activate the simulator by a set of random genes as by pre-existing iterations that exist prior to the simulation run.

Overexpression scheme is based on multiplication of the available miRNA amount by 7 increasing factors (from x1 to x1000). In case the miRNA had not been detected in the naïve cell, an arbitrary starting minimal amount of 0.02% (equivalent of 10 molecule/ cells) are added to the naïve cell (x1).

#### 2.2.2 Analytical methods

Statistical values are that are based on correlations were performed using standard Phyton statistical package. For annotation enrichment statistics and visualization Enrich (Kuleshov et al., 2016) was used. Clustering was performed by the k-mean classification. We used the unsupervised Elbow method to test the consistency within clusters by the percentage of variance explained. (i.e. the ratio of the between-group variance to the total variance). A change in the slop is indicative to the optimal number of clusters in that dataset. Standard statistical tests were applied to provide p-value for protein set comparisons.

## 3 RESULTS

### 3.1 Assessment of the probabilistic approach

The goal of this study is to model the outcome of the miRNA-mRNA network under simplified conditions of translational arrest, mimicking the stochastic nature of miRNA regulation in living cells. Classifying genes into different sets reduces the complexity of the analyzed system. Moreover, it provides new insights on genes that are shared by their sensitivity to miRNA regulation.

The nature and extent of miRNA regulation in living cells is depicted by the absolute quantities, composition and stoichiometry of the main players of the network, i.e., the miRNAs and mRNAs (Arvey et al., 2010). Evidently, the molecular interactions of miRNA and mRNA within a cell is a stochastic process. The specific composition in cells, and binding probability dictate the effectiveness of attenuation of gene expression. Systematic analysis of the miRNA-mRNA interaction network shows that the miRNA regulation operates under tight stoichiometric constrains in living cells. Data from HeLa for miRNAs and mRNAs are extracted from repeated NGS experiments. Total of 539 miRNA types were mapped and 16,236 expressed mRNAs (not including miRNAs). We developed COMICS (Competition of miRNAs Interactions in Cell Systems) as an iterative simulator which was designed to capture quantitative considerations of miRNA-mRNA interaction in living cells. Fig. 1 illustrates the scheme from a cellular perspective while focusing on the probabilistic framework.

**Figure 1:**
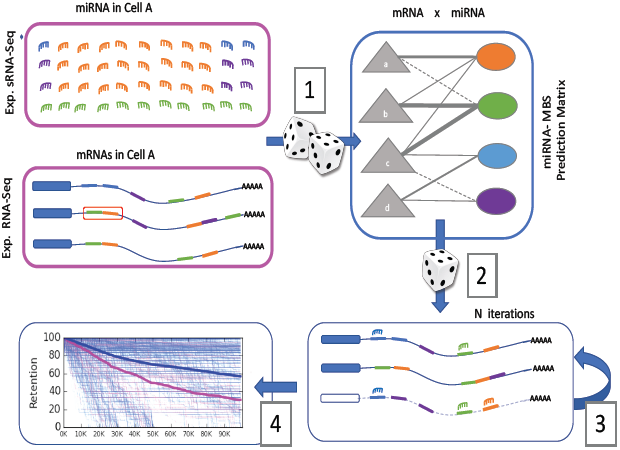
Schematic procedure of the probabilistic nature of COMICS.

### 3.2 COMICS performance

COMICS iterations capture the stochastic process that occurs in cells according to the quantities and the ratio of miRNA to mRNAs. The sampling process (Fig. 1, [1]) driven by the distribution miRNAs and mRNAs according to experimental measurements (Fig 1, pink frames). Each mRNA is characterized by the types and positioning of its miRNA binding sites (MBS) at the 3’-UTS of the transcript. The interaction prediction table is associated with a probability-based scores for any specific pairs of miRNA and MBS in the context of a specific mRNA. Recall that the expression profiles of miRNAs and mRNAs are cell-type specific.

In each iteration, a miRNA is sampled randomly, according to the cell’s miRNA abundance and composition. Next, one of its target gene is chosen randomly according to the measured expressed mRNAs distribution. In the following stochastic step, the randomly chosen miRNA and its target are expected to interact according to the probability of such pairing to be successful. The binding probabilities are based on the miRNA-target prediction of TargetScan (Fig. 1 [2]). It provides a sparse table of miRNA-MBS interactions and reports on 1.2M pairs (see Methods). Each miRNA-MBS interaction is associated with a probabilistic score that is a proxy for the level of confidence for that interaction, and can be considered the probability of effective binding for any specific pair. Following a successful binding event, the distribution of the miRNAs and the mRNAs is updated accordingly (Fig. 1 [3]).

Following a successful pairing, the status of the mRNA is changed (i.e., ready to degradation), and it is marked as ‘occupied’. Upon binding, it may be engaged in additional binding of miRNAs. For MBS that are in a physical proximity to each other, the overlapped interaction is being excluded. The occupied mRNA is marked for degradation with some delay (e.g., 1000 iteration runs) that mimics the likely instance of a cooperative binding on a target by multiple miRNAs prior to its degradation. Once the occupied mRNA is removed from the system, all miRNAs that were bound to it are relieved to the free miRNA pool. As a result, the stoichiometry of miRNA to mRNA is gradually changing with an increase in the apparent ratio of miRNAs to free mRNAs in the cell.

Results of such simulation is illustrated in Fig. 1[4]. In this case, hsa-mir-155 was overexpressed. The decay rate for all genes is shown. The target and non-target genes are differently coloured (pink for targets, blue for non-targets). Cell state is defined as the retention levels (%) of the unbounded genes at the end of the simulation run. COMICS routinely runs for 100,000 iterations.

COMICS supports a wide set of configurable parameters (see Methods), and was tested against the paired interactions as presented by CLASH (Helwak et al., 2013). By including the input from the cells (cell line HEK293) we validated that the overlap with CLASH results is statistically significant (10k to 100k iterations, stable p-value <0.002) (Mahlab-Aviv, 2018). These tests show consistent overlap of the experimental and computational schemes. Thus, confirming the relevance of the simulator protocol to successfully mimic experimental results. In this study, we operating the system for a ‘transcriptional arrest’ mode.

### 3.3 Dynamic gene classes

K-means classification was performed on all genes according to their retention profiles throughout one simulation run (using the Elbow method for 100k COMICS iterations).

Fig. 2A presents the clustering result for k=5 (ensuring a minimal class size). The average behaviour of the dynamic classes of all genes is shown (total 750 genes). The number of genes associated with the different dynamic class are very skewed, with cluster #1 occupies 69% of the genes and <4% are associated with the fast decay cluster (cluster #5). Inspecting the decay rate of the low retention clusters (cluster #4 and #5, 7% of total genes) shows that while the rates are very different the endpoint converged to a similar low retention level.

**Figure 2:**
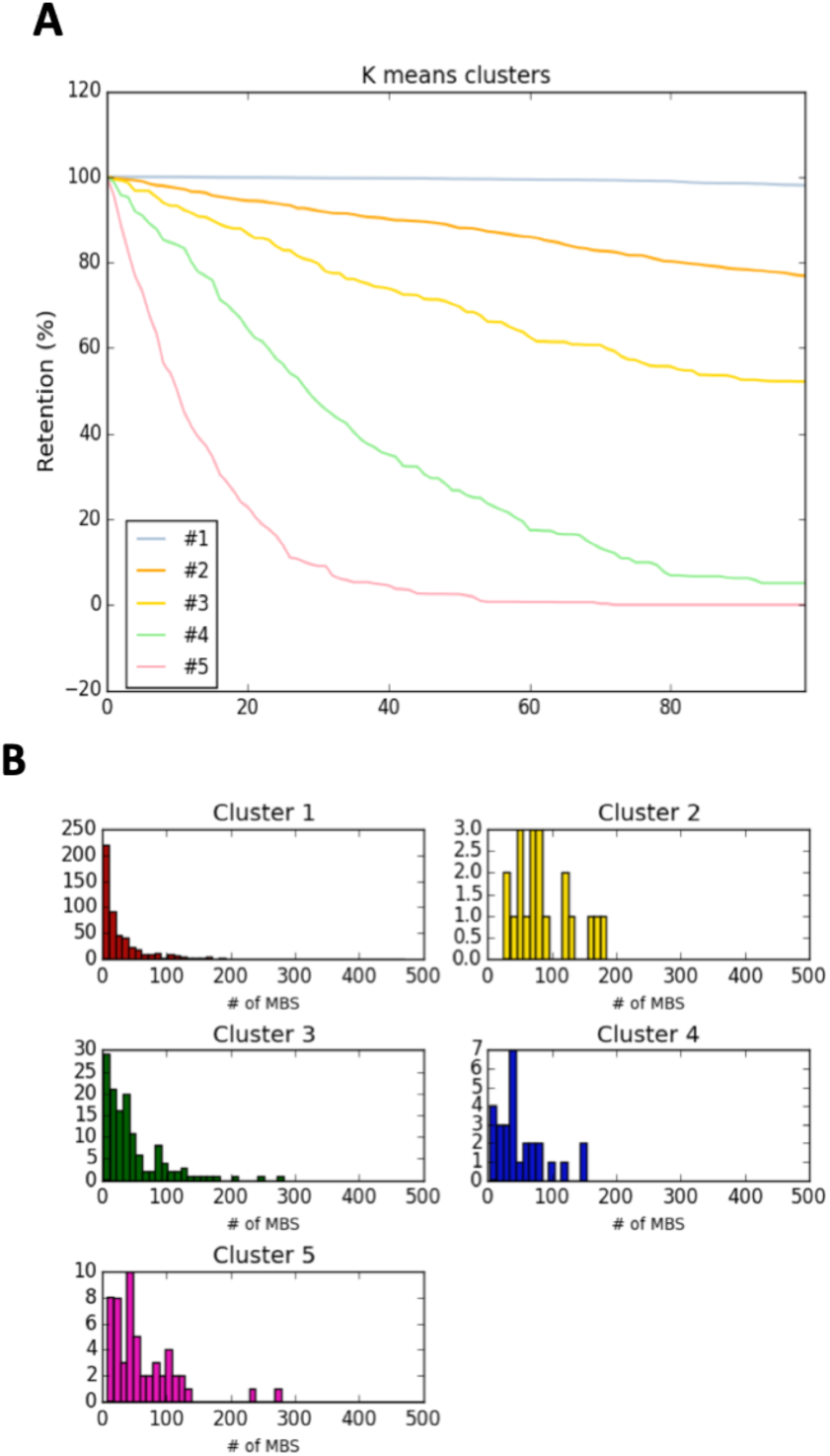
Dynamic classification and analysis. (A) Dynamic classes of Hela cells along 100k iterations of a single simulation run. (B) The distribution of number of MBS per gene in the 5 dynamic classes.

Cluster #5, the lowest retention rate, contains genes that are enriched in annotation of transcription regulators and splicing factors. On the other hand, cluster #1, the most stable one, contains genes of the translation machinery, ribosomal subunits, chaperones, and cytoskeletal components. Genes that are associate with specific functional groups provide additional support for the benefit of classifying genes for reducing the system’s complexity.

Fig. 2B shows a wide distribution of the number of MBS per gene for clusters #2-#5 (irrespectively to their size), with the exception of cluster #1. The number of MBS in the mRNAs in cluster #1 is skewed towards low number of MBS, with an average of 29.7 MBS per gene, while for the rest of the clusters the average is 54.1 MBS per gene. The statistical analysis on average amount of molecules in the cells, length of 3’-UTR are insignificantly different among all dynamic classes. We conclude that class #1 is characterized by relatively low potential to be engaged in successful interactions due to the low binding potential of its genes.

#### 3.3.1 Exhaustive perturbations of miRNAs

To determine the underlying partition between properties that can be attribute to the specific cell type (i.e. profile of its miRNA and mRNAs) and features that are evolutionary driven (i.e. number of MBS in the 3’-UTR, length of 3’-UTR), we activated a systematic analysis on hundreds of cell states derived from a broad range of manipulations of all expressed miRNAs. We applied COMICS simulations using overexpression paradigm in HeLa cells for 248 miRNAs that were compiled according to their representation in the miRNA-MBS TargetScan prediction table (see Methods). We multiply the basal abundance (x1) of each of these miRNA families by the following factors: x3, x9, x18, x90, x300 and x1000. For each such factor (f), a final retention table was computed, and the cell states in the end of the COMICS run were monitored.

Fig. 3 shows the pattern of the retention (%) for a matrix of miRNAs (columns) and genes (same gene list, Fig. 2). The 4 panels are associated with Mfij at factors x3, x9, x90 and x1000. A cell state results are presented by a color-coded table with 750 genes (rows) whose initial expression level exceeds a pre-determined threshold, and 248 miRNAs (columns, organized alphabetically). Each of the listed miRNA was overexpressed by the marked factor. Therefore, each cell in the matrix Mfij is the final retention of gene i after 100k iterations of COMICS for the overexpressed experiment of miRNA j (Fig 3).

**Figure 3:**
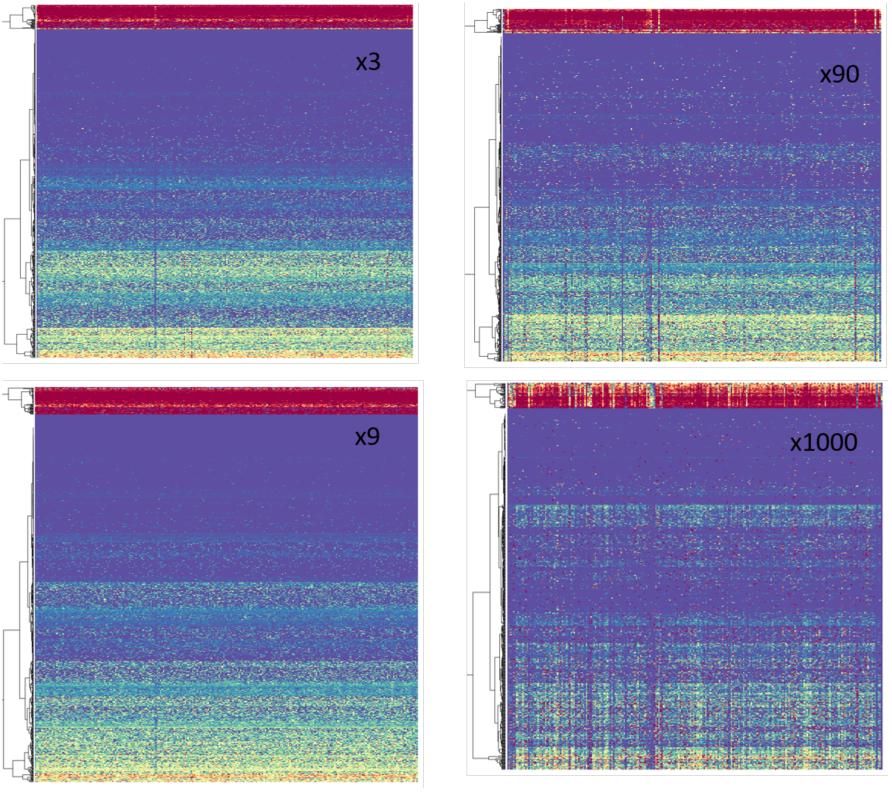
Retention values following several factors of overexpression from HeLa. Rainbow colour code: red=0 to purple=100% retention level at the end of the simulation run. Each panel consists of 248 miRNAs (columns) and 750 genes (rows). The overexpression factors are indicated (x3 x9, x90, x1000). The matrices are clustered by genes.

Inspecting the Mfij (4 factors of overexpression conditions) reveals that genes are naturally clustered by their final retention level. For example, the top ‘red’ rows represent a set of genes that were very sensitive (red=0% retention; purple=100% retention) to almost any manipulations. Clear observations can be drawn from inspecting these matrices (Fig. 3): (i) Large number of coordinated behaviours is evident for each set of genes. It is reflected by observing a similar colour by the entire row across most miRNA columns. A large number of rows displays this distinctive property. (ii) As the overexpression factor increases (i.e. x1000) the pattern of the columns (i.e. specific miRNAs) becomes evident. It is reflected by observing a similar colour across the entire columns, and across most rows.

#### 3.2.2 Perturbation pair ratio classes

In view of the observation that many genes behave similarly with respect to their retention levels at a wide range of Mfij, we tested whether genes can be characterizing by their degree of sensitivity to the abundance of miRNAs (as studied by exhaustive overexpression tests).

Following the relative changes of each gene retention in each miRNA, and in each overexpression factor, we computed the retention ratio between any tested overexpression factors. Formally, we computed the value of Mfij/ Mkij, which is the ratio of the retention of a specific gene (simulation at 100k iterations) in a specific miRNA overexpression of factor f, and its retention in the same miRNA overexpression factor k (Fig. 4A). For visualization purpose, a discretization was applied for which ratio that is >2 folds. It implies that the retention of genes i in the overexpression of miRNA j by factor f is higher than its retention where miRNA j was overexpressed by factor k (Fig 4B, blue cells). However, ratio that is <0.5 implies that in factor f the gene is more prone to degradation with respect to factor k (Fig 4B, red cells).

**Figure 4:**
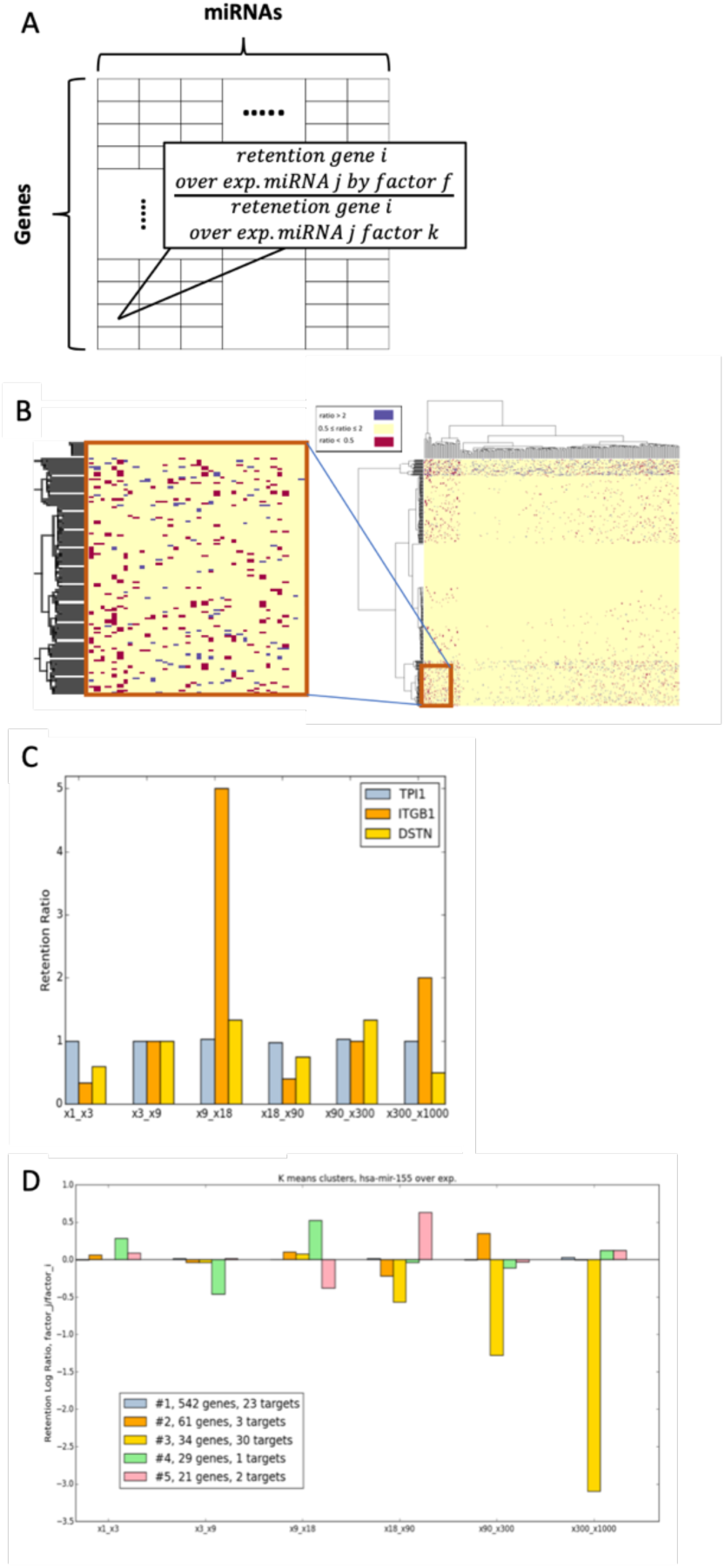
Overexpression ratio (OXR) classes. (A) Scheme for the presented Mfij/ Mkij ratio retention matrix. (B) Ratio matrix for x90/x9 and a zoon-in for a small section. Colours only marked genes that show a significant ratio difference (by the predetermined differential folds of >2, and <0.5). (C) Examples of 3 genes according to the retention ratio from 6 ratio-matrices. (D) OXR-clusters measuring the genes associated with five OXR-classes by their sensitivity to hsa-mir-155 manipulations, represented for 6 ratio matrices (on x-axis). The numbers of expressed targets of hsa-mir-155 that are included in each OXR-class are indicated in the Figure legend).

Fig 4C illustrates the retention ratios of selected three genes (for illustrative purposes). It is shown that following overexpression of hsa-mir-155 the gene TPI1 remains stable throughout any of the tested retention ratios. As expected, TPI1 belongs to the dynamic class of genes that are extremely stable in the system. A different behaviour is observed for ITGB1 whose expression is very unstable and sensitive to a minor change in overexpression factors (e.g. x18/x9). The non-monotonic behaviour of ITGB1 and DSTN are evident.

Fig 4D illustrates the gene sensitivity as measured by the retention rate of different genes in the case of hsa-mir-155 for 6 different pairs of factors: (x1, x3), (x3, x9), (x9, x18), (x18, x90), (x90, x300), and (x300, x1000). Same analysis (as in Fig. 4C) was performed for all expressed genes. The results for all genes for a specific miRNA overexpression were clustered by K-means clustering algorithm (a cluster must contain >5 genes; OXR-classes). The analysis reveals that OXR-classes display different sensitivity pattern with respect to the pair-overexpression retention ratio.

Fig. 4D shows the partition of all genes to 5 clusters (marked #1 to #5). The majority of the genes (∼71%, cluster #1, coloured grey) are indifferent to the levels of overexpression factors. However, the rest (∼29%) of the genes are sensitive to some extend for the overexpression factors that was used. For example, cluster #3 (Fig 4D, yellow) contains genes that their retention rate is drastically decreased by OXR, as the overexpression factor increases (f). It is satisfying to note that most hsa-mir-155 expressed target genes (52%) belong to cluster #3. However, other target genes are associated with additional clusters with most of them belong to cluster #1.

The behaviour of 4 selected miRNAs according to their OXR-classes, for 6 matrix ratio combinations is shown (Fig. 4D and Fig. 5). The represented miRNAs are expressed at different order of magnitudes. miRNAs that are highly expressed (e.g., hsa-mir-7, 4.2%) were analyzed as well as miRNAs that are expressed at a low level (hsa-mir-320, 0.15%; hsa-mir-155, 0.02%).

**Figure 5:**
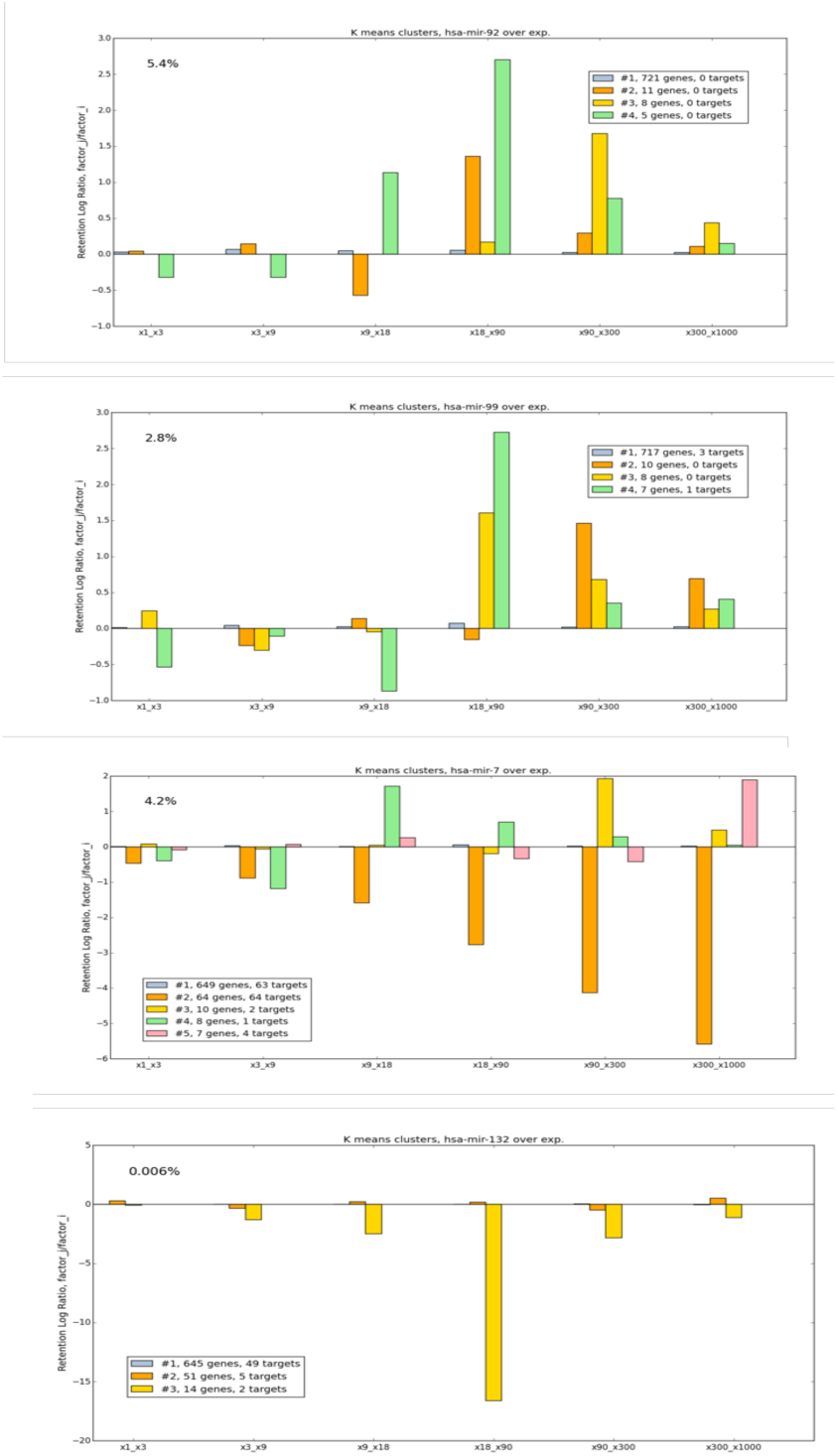
Example of OXR classes for different miRNAs, following manipulations of hsa-mir-7, hsa-mir-99, has-mir-24, and hsa-mir-320. The numbers of the targets of each tested miRNA are indicated (see color legend). miRNA expression (%) is marked. The values at y-axis are expressed by log2.

Fig. 5 shows several behaviours associated with the OXR-classes: (i) In almost all instances, OXR-class that includes most targets of the subjected miRNA, decreases monotonically with a maximal effect seen for ratio of the highest overexpression pair (i.e. x1000/x300). This observation is generalized, irrespectively to the initial expression level of the miRNAs. This strongest effect is associated with OXR-classes that include only targets (e.g., cluster #2 in hsa-mir-7). (ii) Some OXR-classes show a characteristic behaviour that cannot be trivially anticipated. For example, a switch in the trend of cluster #4 of hsa-mir-99 shows that for the ratio of x18 and x90 the effect of retention does not follow the behaviour at any of the other expression ratio pairs. (iii) For some miRNAs there are no target detected in the list of the analyzed genes (e.g., hsa-mir-92). In such instance, we do not detect a ‘target’ OXR-class and the trend of the OXR-classes is more indicative for a gene stabilized pattern (e.g., hsa-mir-99 and hsa-mir-92). (iv) For all miRNAs, the largest OXR-classes includes 82% to 91% of the analyzed genes. This general observation implies that most genes are insensitive to the perturbation according to the pair-ratio. (v) Some of the clusters show extreme increase or decrease in the retention rates (e.g., hsa-mir-320 for cluster #3, at x90/x18). This implies the high sensitivity of these gene sets to a specific miRNA abundance.

We illustrated OXR-classes for 6 ratio-matrices. The comparison and analyses are valid for each miRNA (total 248), and for 21 pairs the 7 overexpression factors tested.

## 3 DISCUSSION

Cells’ behavior cannot be extracted from direct measurement of the composition of miRNAs and mRNAs (Arvey et al., 2010; Landgraf et al., 2007). Most insights on regulation of gene expression by miRNAs in the complexity of the cells are based on CLIP-Seq and CLASH methodologies (Li et al., 2014). Based on many such studies, it was concluded that detailed quantitative considerations of miRNA and mRNA govern the dynamics and the steady state of gene expressed in cells (Bosson et al., 2014; Hausser and Zavolan, 2014). Nevertheless, the underlying rules for post-transcriptional regulation by miRNAs are still fragmented (Erhard et al., 2014).

In all the presented COMICS results we consider miRNA pool to be constrained by the amount of AGOs in the cells. Several studies estimate in each cell ∼50k AGO molecules (but huge variation is reported). COSMIC is insensitive to such quantitation debates as the sampling protocol (Fig 1) relies on probabilistic formulation. Under varying levels of miRNA overexpression, the loading of miRNA molecule on AGO is driven by the actual distribution of that miRNAs.

The OXR-classes aim to capture the system dynamics rather than the dynamics of the gene expression downregulation by miRNAs. We were able to cluster genes to their OXR-classes by performing hundreds of simulations allowing a robust assessment of cell states. For most instances, under all conditions, the majority or the expressed genes are not sensitive to the matrix-ratio measures. Namely, the final retention that is achieved in all conditions of overexpressed miRNAs is unchanged (Fig. 5, y-axis =0). In a smaller set of genes, miRNAs strongly regulate their targets when a switch in the abundance occurs from one level to another.

In this study we consider two sets of gene classes: dynamic-class (Fig. 2) and OXR-classes (Figs 4-5). These two complementary types of classes capture different aspects of miRNA regulation dynamics. Results from the dynamic-class show than genes that are likely to be successfully targeted are those having large number of MBS at the 3’-UTR (Fig. 2). However, the dynamic-classes #2 to #5 are not distinguishable by such feature. Specifically, cluster #2, #3, #4, #5 are associated with 50.3, 71.6, 63.3 and 47.6 average MBS per gene. In the future, and using analysis from different additional cell lines, we will extend the analysis to include evolutionary based features such as the distribution of MBS along the 3’-UTR, the number of alternative spliced variants that affect miRNA regulation, the presence of MBS that binds any of the most abundant miRNAs in the cell. Addition of cell specific features to the clustering protocol (i.e. cell specific genes variants) will benefit the refinement of the classification protocol.

It was shown that miRNA profile is carefully regulated to promote and stabilize cell fate choices (Shenoy and Blelloch, 2014). In accord with this notion, we show a general trend for the highly expressed miRNAs. Actually, many large-scale miRNA overexpression experiments (e.g., (Lim et al., 2005; Wang and Wang, 2006)) overlooked and ignored the gene sensitivity to the actual amounts of miRNAs. For the majority of studies, the degree of overexpression of miRNA is not even reported. We suggest that it may contribute to the inconsistency that is often observed among experimental results. Using COMICS dynamic comparison benefits an accurate prediction considering the exact levels of miRNA overexpression.

Despite over two decades of research in the miRNA field, basic principles remained to be discovered. Current miRNA-mRNA prediction tools suffer from a large number of false positive. The experimental methodologies (e.g. CLASH and CLIP-Seq) that are based on capturing the interactions followed by sequencing are not always consistent, show poorly reproducibility (Lu and Leslie, 2016). We suggest that a representation of genes by their dynamic and OXR-classes can yield an accurate yet simplistic model for miRNA regulation. This study illustrates the use of COMICS results for gene classification. It further stresses the importance of quantitative view for miRNA regulation modelling.

In pathological cells, such as in cancer, a quantitative change in the amounts of miRNAs is often the most significant molecular change observed in early phase of cancer development. Assessing changes in behaviour of representative genes from OXR-classes could benefit cancer diagnosis. Some OXR-classes may serve as indicator for a shift in cell states and identity. The available of a very small number of coherent gene classes, argues that cells display an unexpected robustness with respect to the miRNA regulation. The ability to classify genes according to dynamic overlooked features carries its potential to improve cell modelling, and improving the understanding of cellular miRNA regulation.

## ACKNOWLEDGEMENTS

The study was supported in part by ERC grant 339096 on High Dimensional Combinatorics (to SMA). We benefit from the core tools and resources supported by ELIXIR. The study is partially supported by funds # 9660/2019 from Yad Hanadiv (to ML).

## Notes

### Competing Interest Statement

The authors have declared no competing interest.

## REFERENCES

Arvey, A., Larsson, E., Sander, C., Leslie, C.S., and Marks, D.S. (2010). Target mRNA abundance dilutes microRNA and siRNA activity. Mol Syst Biol 6, 363.

Balaga, O., Friedman, Y., and Linial, M. (2012). Toward a combinatorial nature of microRNA regulation in human cells. Nucleic Acids Res 40, 9404–9416.

Bertoli, G., Cava, C., and Castiglioni, I. (2015). MicroRNAs: New Biomarkers for Diagnosis, Prognosis, Therapy Prediction and Therapeutic Tools for Breast Cancer. Theranostics 5, 1122–1143.

Bosson, A.D., Zamudio, J.R., and Sharp, P.A. (2014). Endogenous miRNA and target concentrations determine susceptibility to potential ceRNA competition. Molecular cell 56, 347–359.

Chekulaeva, M., and Filipowicz, W. (2009). Mechanisms of miRNA-mediated post-transcriptional regulation in animal cells. Curr Opin Cell Biol 21, 452–460.

Eichhorn, S.W., Guo, H., McGeary, S.E., Rodriguez-Mias, R.A., Shin, C., Baek, D., Hsu, S.H., Ghoshal, K., Villen, J., and Bartel, D.P. (2014). mRNA destabilization is the dominant effect of mammalian microRNAs by the time substantial repression ensues. Mol Cell 56, 104–115.

Erhard, F., Haas, J., Lieber, D., Malterer, G., Jaskiewicz, L., Zavolan, M., Dolken, L., and Zimmer, R. (2014). Widespread context dependency of microRNA-mediated regulation. Genome Res 24, 906–919.

Hausser, J., and Zavolan, M. (2014). Identification and consequences of miRNA-target interactions-- beyond repression of gene expression. Nature Reviews Genetics 15, 599.

Kozomara, A., and Griffiths-Jones, S. (2013). miRBase: annotating high confidence microRNAs using deep sequencing data. Nucleic acids research 42, D68–D73.

Landgraf, P., Rusu, M., Sheridan, R., Sewer, A., Iovino, N., Aravin, A., Pfeffer, S., Rice, A., Kamphorst, A.O., and Landthaler, M. (2007). A mammalian microRNA expression atlas based on small RNA library sequencing. Cell 129, 1401–1414.

Li, J.H., Liu, S., Zhou, H., Qu, L.H., and Yang, J.H. (2014). starBase v2.0: decoding miRNA-ceRNA, miRNA-ncRNA and protein-RNA interaction networks from large-scale CLIP-Seq data. Nucleic Acids Res 42, D92–97.

Lim, L.P., Lau, N.C., Garrett-Engele, P., Grimson, A., Schelter, J.M., Castle, J., Bartel, D.P., Linsley, P.S., and Johnson, J.M. (2005). Microarray analysis shows that some microRNAs downregulate large numbers of target mRNAs. Nature 433, 769–773.

Lu, J., Getz, G., Miska, E.A., Alvarez-Saavedra, E., Lamb, J., Peck, D., Sweet-Cordero, A., Ebert, B.L., Mak, R.H., Ferrando, A.A., et al. (2005). MicroRNA expression profiles classify human cancers. Nature 435, 834–838.

Lu, Y., and Leslie, C.S. (2016). Learning to Predict miRNA-mRNA Interactions from AGO CLIP Sequencing and CLASH Data. PLoS Comput Biol 12, e1005026.

Moore, M.J., Scheel, T.K., Luna, J.M., Park, C.Y., Fak, J.J., Nishiuchi, E., Rice, C.M., and Darnell, R.B. (2015). miRNA-target chimeras reveal miRNA 3’- end pairing as a major determinant of Argonaute target specificity. Nat Commun 6, 8864.

Pelaez, N., and Carthew, R.W. (2012). Biological robustness and the role of microRNAs: a network perspective. Curr Top Dev Biol 99, 237–255.

Peterson, S.M., Thompson, J.A., Ufkin, M.L., Sathyanarayana, P., Liaw, L., and Congdon, C.B. (2014). Common features of microRNA target prediction tools. Frontiers in genetics 5.

Pinzon, N., Li, B., Martinez, L., Sergeeva, A., Presumey, J., Apparailly, F., and Seitz, H. (2017). microRNA target prediction programs predict many false positives. Genome Res 27, 234–245.

Rajewsky, N. (2006). microRNA target predictions in animals. Nature genetics 38, S8.

Shenoy, A., and Blelloch, R.H. (2014). Regulation of microRNA function in somatic stem cell proliferation and differentiation. Nature Reviews Molecular Cell Biology 15, 565.

Wang, X., and Wang, X. (2006). Systematic identification of microRNA functions by combining target prediction and expression profiling. Nucleic Acids Res 34, 1646–1652.

Zhang, Y., Wei, W., Cheng, N., Wang, K., Li, B., Jiang, X., and Sun, S. (2012). Hepatitis C virus-induced up-regulation of microRNA-155 promotes hepatocarcinogenesis by activating Wnt signaling. Hepatology 56, 1631–1640.

